# Multiple Conformations of Gal3 Protein Drive the Galactose Induced Allosteric Activation of the *GAL* Genetic Switch of *Saccharomyces cerevisiae*

**DOI:** 10.1101/032136

**Authors:** Rajesh Kumar Kar, Hungyo Kharerin, Ranjith Padinhateeri, Paike Jayadeva Bhat

**Author notes:** has performed all the experimental work (Rajesh Kumar Kar). has performed all the molecular dynamics simulations work (Hungyo Kharerin). Current Address: Department of Biochemistry, College of Agricultural and Life Sciences, 433 Babcock Drive, University of Wisconsin-Madison, WI 53706.

## Abstract

Gal3p is an allosteric monomeric protein which activates the *GAL* genetic switch of *Saccharomyces cerevisiae* in response to galactose. Expression of constitutive mutant of Gal3p or over-expression of wild-type Gal3p activates the *GAL* switch in the absence of galactose. These data suggest that Gal3p exists as an ensemble of active and inactive conformations. Structural data has indicated that Gal3p exists in open (inactive) and closed (active) conformations. However, mutant of Gal3p that predominantly exists in inactive conformation and yet capable of responding to galactose has not been isolated. To understand the mechanism of allosteric transition, we have isolated a triple mutant of Gal3p with V273I, T404A and N450D substitutions which upon over-expression fails to activate the *GAL* switch on its own, but activates the switch in response to galactose. Over-expression of Gal3p mutants with single or double mutations in any of the three combinations failed to exhibit the behavior of the triple mutant. Molecular dynamics analysis of the wild-type and the triple mutant along with two previously reported constitutive mutants suggests that the wild-type Gal3p may also exist in super-open conformation. Further, our results suggest that the dynamics of residue F237 situated in the hydrophobic pocket located in the hinge region drives the transition between different conformations. Based on our study and what is known in human glucokinase, we suggest that the above mechanism could be a general theme in causing the allosteric transition.

**Abbreviations**
d.o.: drop out
2-DG: 2-deoxy-galactose
Gal3p: Gal3 protein
EtBr: Ethidium bromide
GAL3^c^: GAL3 constitutive mutant
ORF: open reading frame
EPPCR: Error Prone PCR
CMD: Canonical molecular dynamics
TMD: Targeted molecular dynamics
SO: Super-open

## Introduction

Information transfer through molecular recognition plays a vital role in cellular processes such as catalysis, transcriptional regulation and differentiation. This occurs mainly through allostery, wherein ligand binding at one site induces a conformational change at another site (Monod *et al*. 1965; James & Tawfik 2003). This alteration in the conformation of the protein is conveyed to the downstream target in a highly calibrated and quantitative fashion. Allostery was originally thought to occur only in multimeric proteins either through the induced fit or through a conformational selection mechanism as proposed in KNF (Koshland 1958) and MWC (Monod *et al*. 1965) models respectively. Subsequent success of the structural studies in exploring the MWC model suggested that allostery in general can be understood from a structural standpoint (Marsh & Teichmann 2015). A model for explaining allostery without invoking a conformational alteration was proposed based on thermodynamic considerations (Cooper & Dryden 1984). While the above models provide a fundamental conceptual framework for our current understanding, the allosteric behaviour of many proteins need not necessarily follow one or the other mechanisms nor the postulates of the above models are adequate to explain the complex allosteric behaviour of many proteins. For example, it has been suggested that strong and long-range protein-ligand interactions favour induced fit model while weak and short range interactions favour conformational selection mechanism (Okazaki & Takada 2008). On the other hand, the rate of conformational transition could determine whether induced fit or the conformational selection mechanism dominates (Hammes *et al*. 2009; Vogt & Di 2012). Recently, computational analysis has indicated both conformational selection and induced fit mechanisms operate during the allosteric transition of LAO proteins (Silva *et al*. 2011). In contrast to the above studies, how a ligand acts as an agonist as well as an antagonist of the same protein is difficult to explain using the KNF or MWC model (James *et al*. 2003; Motlagh & Hilser 2012).

As opposed to the original postulate of MWC model wherein allosteric protein visits only a few allowed conformations, it was suggested quite early on that allosteric proteins are likely to visit a large number of conformations (Weber 1972). It has been suggested that allostery is manifested due to the “redistribution of the conformational ensemble” in the presence of ligands and therefore is an intrinsic feature of all dynamic proteins (Gunasekaran *et al*. 2004; Boehr *et al*. 2009; Blacklock & Verkhivker 2014). A large body of experimental evidence supports this possibility of population shift (or conformational selection) induced by ligands. For example, Nitrogen regulatory protein C (NtrC) is a signalling protein involved in the transcriptional regulation of genes responsible for nitrogen assimilations in bacteria and its activation occurs through a phosphorylation event (Kern *et al*. 1999). Using NMR, it has been demonstrated that this protein exists as an equilibrium mixture of active and inactive states, indicating that phosphorylation only shifts the equilibrium to the active state (Volkman *et al*. 2001). A similar process of phosphorylation-mediated active-inactive conformational shift is also observed in CheY, a monomeric single-domain response regulator protein involved in chemotaxis (Schuster *et al*. 2001). Recent evidence suggest that CheY does not undergo a two-stage concerted mechanism, rather employs a combination of the KNF and MWC mechanisms to function like a switch (McDonald *et al*. 2012). In contrast to CheY, human glucokinase, a monomeric protein with two distinct domains, has been shown to pre-exists in closed state prior to the introduction of ligands, even though this state may be the minor species (Kamata *et al*. 2004; Zhang *et al*. 2006). Thus, although the induced fit mechanism is necessary, the conformational selection appears to be the driving force for the allosteric transition.

Thus, at the conceptual level, thermodynamic, population shift, and structural basis of allostery may appear disparate, but at the fundamental level, allostery needs to be explained from all the above perspectives and accordingly a unified view of allosetry has been proposed (Tsai & Nussinov 2014). Independent of the above, it has been proposed that few key residues are associated with structural stabilization, allosteric couplings and correlated motions (Lockless & Ranganathan 1999; Sinha & Nussinov 2001). Literature is replete with examples, wherein, substitution of single key amino acid residues results in the stabilization of the active conformation (or destabilization of the inactive conformation), which is otherwise attained only after interacting with ligands (Kjelsberg *et al*. 1992; Blank *et al*. 1997). For example, in hamster α-1_b_-adrenoceptor, substitution of Alanine at position 293 to any other residue results in the constitutive activity (Kjelsberg *et al*. 1992). If mutations can stabilize the active conformation, then, in principle, it should be possible to isolate mutations that stabilize the inactive state, without altering the ability of the mutant protein to respond to its ligands, provided conformational shift mechanism is an integral mechanism of allostery. To the best of our knowledge, this corollary has not been experimentally verified. This is probably because, protein in inactive conformations would normally be considered devoid of biochemical activity and unlikely to be analysed further.

In *S. cerevisiae*, galactose activates the allosteric signal transducer Gal3p, which in turn inactivates the repressor Gal80p, thereby activating the Gal4p dependent transcriptional activation of the switch (**Figure S1**). The structure of Gal3p in apo as well as in holo (in presence of galactose and ATP and Gal80p) forms has been solved at a resolution of 2.1 Å (Lavy *et al*. 2012). In addition, an intermediate structure was observed during crystallization only when the concentrations of the ligands used was low. Much before the structure was determined, it was observed that, over-expression of Gal3p activated the *GAL* genetic switch even in the absence of the inducer galactose (**Figure S2C**) (Bhat & Hopper 1992). Based on this, it was hypothesised that Gal3p exists in active and inactive conformations and galactose merely increases the active form of Gal3p (**Figure S2**). This hypothesis was subsequently supported by isolating mis-sense mutations, which confer Gal3p, the ability to activate the *GAL* switch in the absence of galactose (**Figure S2B**) (Blank *et al*. 1997). If the above hypothesis is true, then it should be possible to isolate mutations that stabilize the protein in inactive conformation and yet such a mutant be able to respond to galactose.

In this study, we report the isolation of a novel Gal3p mutant with V273I, T404A and N450D substitutions, using a dual screening strategy. The ability of this Gal3p mutant to activate *GAL* switch upon over-expression is very less as compared to the wild-type, but activates the *GAL* switch in response to galactose. Genetic studies indicated that all the three substitutions are essential for conferring the phenotype described above. Molecular dynamic studies, which showed that unlike the wild-type Gal3p, the mutant form has higher propensity to exist in super-open conformation. We discuss the implications of these observations in the context of glucokinase, an allosteric protein, which has many functional overlaps with Gal3p and both seem to follow a similar pathway of allosteric transition.

## Materials and Methods

### Yeast strains, media and growth conditions

Yeast strains (**Table S1**) were grown either in yeast extract, peptone and dextrose (YEPD) or in synthetic medium supplemented with 3% (v/v) glycerol plus 2% (v/v) potassium lactate (gly/lac), 2% glucose or 2% galactose (sigma, Germany) as carbon source. Whenever required, Ethidium bromide (EtBr) and 3-amino-1, 2, 4-triazole (3-AT) (sigma, Germany) was added to the medium to a final concentration of 20μg/ml and 10mM respectively. Yeast transformation was done by lithium acetate mediated protocol (Gietz & Woods 2002). *E. coli* transformation, plasmid amplification and isolation was carried out as with standard protocols (Sambrook 1989).

## Plasmid and Strain construction

### Construction of *PGAL1GFP* (*pYIPLac211GAL1GFP*) *and pRKK29* (*pYIPLac204GAL1GFP*)

A 1.5 kb cassette containing *P_GAL1_GFP* isolated from *YCpGAL1-GFP* (Stagoj *et al*. 2005) as a *Pst*I*-Eco*RI fragment was ligated at the corresponding sites of *YIpLac211* to obtain *pGAL1GFP* (*pYIpLac211GAL1GFP*) as described before (Kar *et al*. 2014). The same *P*stI-*Eco*RI fragment was ligated at the corresponding sites of *YIpLac204* to obtain *pRKK29* (*pYIpLac204GAL1GFP*) (**Table S2**).

## Strain construction

### *GAL3* disruption by *KanMx4* cassette

*GAL3* is disrupted in *Sc723* (Blank *et al*. 1997) background by integrating *KanMX4* cassette as described before. *ScRKK6* was generated by disrupting *GAL3* with *gal3KanMX4* cassette in *Sc723* (Kar *et al*. 2014).

### *GAL1* disruption by *KanMx4* cassette

*GAL1* was disrupted by integrating *KanMX4. KanMX4* cassette was obtained by amplifying genomic DNA of *BY4742gal1:Kan* by using primers PJB417 (5’GGATGGACGCAAAGAAGTTTAAT AATCATATTACATGGCATTACCACC3’) and PJB418 (5’GTCCTTGTCTAACTTGAAAATTTTTC ATTGATGTCATACGAC3’). This strain was obtained from EUROSCARF, in which *GAL1* ORF is replaced by *KanMX4* cassette. The resulting PCR fragment contains *KanMX4* flanked by 561bp from upstream and 537bp from downstream of *GAL1* ORF. *Sc385* (*gal3:LEU2*) and *Sc756* (*gal1:URA3,gal3:LEU2*) strains were transformed with this PCR product and transformants were selected in YPD plate supplemented with 200µg/ml of G418 and gives rise to *ScRKK5* and *ScRKK1* respectively. The putative transformants were confirmed by phenotype analysis and PCR by primers PJB428 (5’CCA GACCTTTTCGGTCACAC3’) and PJB427, which is designed 826bp upstream of *GAL1* ORF and 605bp internal of *KanMX4*.

### Construction of *ScRKK7* and *ScRKK39*

*pGAL1GFP* as well as *pRKK29* was linearized with *Eco*RV located within the *URA3* and *TRP2* locus respectively. This linearized *pGAL1GFP* plasmid was integrated at the ura3-52 locus of *Sc723* to obtain the *ScRKK7* strain as described before (Kar *et al*. 2014). The linearized *pRKK29* was integrated at the *trp2* locus of *ScRKK1* to obtain *ScRKK39* strain.

### Error Prone PCR [EPPCR]

*GAL3* ORF was cloned as a 1.6kb *Eco*RI and *Hin*dIII fragment into the corresponding site of *pYJM* resulting in *pYJM3* was available in the laboratory (Murthy & Jayadeva 2000). Error Prone PCR [EPPCR] was done by using the classical protocol (Lin-Goerke *et al*. 1997). To introduce random mutations in *GAL3* ORF, EPPCR was carried out using the primers PJB328 (5’GGAAAGCGGGCAGT GAGCG3’) and PJB329 (5’GGCATGCATGTGCTCTGTATG3’) at mutagenic condition (**Table S3 and S4**). Freshly prepared MnCl2 was always used for the mutagenic condition.

### Site directed mutagenesis

Site directed mutagenesis was performed by using “Quick change site-directed mutagenesis protocol (QCM) developed originally by Stratagene. The oligonucleotides used for site directed mutagenesis are listed in (**Table S5**). The mutations were confirmed by primer-walking sequencing analysis from Scigenome as well as from Chromous biotech.

### 2-Deoxy-galactose (2-DG) toxicity assay

Constitutive expression of *GAL1* in various transformants of *gal3*Δ cell was monitored based on their ability to grow in gly/lac + 2-DG. It has been reported that cells constitutive for galactokinase expression would not be able to grow in a media containing 2-DG due to toxicity (Platt 1984). This toxicity is due to the accumulation of 2-DG-1p. Also, it has been reported that 2-DG does not substitute galactose for the signal transducing activity of Gal3p. Cells, which do not produce galactokinase, do not show toxicity and therefore grow when provided with an alternative carbon source glycerol plus lactate (gly/lac) (Kar *et al*. 2014).

### Histidine growth assay

*ScRKK1* (*gal1*Δ*gal3*Δ) cells were used for this assay, which has a *P_GAL1_::HIS3* cassette integrated at the *LYS2* locus. *ScRKK1* (*gal1*Δ*gal3*Δ) cells were pre-grown to mid-log phase in complete medium containing gly/lac as the sole carbon source. Cells were appropriately diluted and plated on to medium containing (a) uracil d.o. gly/lac (b) uracil d.o. gly/lac medium lacking histidine with and without galactose. Medium lacking histidine contained 3-amino-1,2,4-triazole (3-AT) to reduce background growth due to the leaky expression of HIS3 (Blank *et al*. 1997). We have already used this assays to score the cells, which responds to galactose (Kar *et al*. 2014).

### FACS analysis

Cells were grown in complete medium containing gly/lac and re-inoculated into medium containing 2% glucose and 2% of galactose. Cells were harvested at different time intervals, washed with PBS twice, re-suspended in PBS and kept in ice. A minimum of 50,000 cells were analyzed for each data point. Flow cytometry data was collected using BD FACS ARIA-1 flow cytometer with FITC filter for detection of GFP (Kar *et al*. 2014). Data were collected by BD FACS DIVA software. The overlay and data analysis was done by FCS express 4 flow research edition software (De Novo Software, Toronto, Ontario, Canada). At least three sets of independent experiments were performed for FACS analysis. The FCS express 4 flow research edition software overlaid data were tabulated (**Table S6**).

### Electrophoresis procedure

Protein samples for SDS-PAGE were carried out according to the method of Laemmli (Laemmli 1970). 10% gels were used for all experiments. The electrophoresis was carried out at 15 mA per gel.

### Western blot analysis

Yeast cell extract was prepared by harvesting the cells grown in mid-log phase by centrifuging at 5,000 rpm for 5 min and then washed once with cold autoclaved double distilled water. Cells were then resuspended in cold autoclaved Phosphate buffer saline. Protease inhibitor cocktail and PMSF were added to the final concentration of 2.5 mM and 1 mM respectively. Equal amounts of 0.45 mm diameter cold glass beads were added to each microfuge tube and subjected to vortexing for 1 min, followed by incubation on ice for 1 min. This cycle was repeated for 10 times. Cell lysates were centrifuged at 13,000 rpm in a microcentrifuge for 25 min at 4°C. The protein was quantified by BCA kit (Merck). The protein obtained was boiled with gel loading buffer for 10 min and resolved in 10% SDS polyacrylamide gel. The protein was transferred from the SDS gels to nitrocellulose membrane at 130 mA for 3 hour in transfer buffer. The transferred protein was blocked with blocking buffer [PBS pH8.0 containing 3% (w/v) skimmed milk powder and 0.02% (v/v) Tween_20_] for overnight. The blot was incubated for 1 hour in 1:1000 purified antibody for detection of Gal3 protein. The blot was then probed with the secondary antibody conjugated with alkaline phosphatase at 1:5000 dilutions. Finally, blot was developed in developing buffer containing NBT and BCIP. The protein bands were quantified by Image J software (Schneider *et al*. 2012).

### Protein models

The PDB coordinates of Gal3p, chain C of 3v2u (closed) and chain A of 3v5r (open) (Lavy *et al*. 2012) in which any ligand present was removed, were used as the starting structures for CMD simulations in closed and open apo-states respectively. We retained ATP and galactose to simulate the liganded structures. Since we do not have crystal structures for Gal3p mutants, Gal3p^IAD^, Gal3p^D368V^ and Gal3p^F237Y^, we derived their structures by homology modeling with crystal structures of Gal3p as templates. All modeling and fixing of missing residues were carried out using the online version of homology modeler (Sali *et al*. 1995) (http://toolkit.tuebingen.mpg.de/modeller).

### Computational Details

All MD simulations were run with NAMD package (Phillips *et al*. 2005) using CHARMM force fields (Brooks *et al*. 2009) and Tip3p (William L. Jorgensen 1983) water model. VMD (Humphrey *et al*. 1996) “autoionize” package was used to neutralize the whole system. Solvent box size was chosen in such a way that the minimum distance from the protein to the wall is 15Å. This will ensure there is no direct interaction with its own periodic images. Covalent bonds involving hydrogen atom are made rigid by applying SHAKE. We have used CMD to study equilibrium dynamics around native conformations. For all protein systems, energy was minimized using conjugate gradient algorithm first for protein alone, then protein + water and finally protein + water + ions. The solvated systems were heated to 300K and then equilibrated as NPT ensemble in two steps. First, protein was fixed and the whole system was equilibrated for 1200ps. Second, the whole system was equilibrated for 1200ps with no constraint. The MD integration time step used was 2.0 fs. In all the CMD simulations, coordinates were saved at every 1ps interval for the analyses. Cα backbone root mean square deviation (RMSD) of simulation trajectory as a function of time was plotted in the **Figure S3A** and **B** to assess simulation stability and to check structural drift. To give enough relaxations to the modeled residues (missing residues and mutation residues) in 3D environment, we set aside a good 20 ns (shaded region in **Figure S3A** and **B**) as a relaxation step and the remaining last 100 ns trajectory was selected to study equilibrium dynamics of proteins.

To study conformational transitions along activation pathway, we have employed TMD. Here, an extra force given by the derivative of potential energy, *U_T M D_* = 0.5×*k*([RMSD(t) – RMSD_0_ (t)]^2^)/*N*, is added to the existing force-field to drive the protein from open (start) to closed (target) conformation. *N* is the number of targeted atoms to which the force is applied and k is the force constant in kcalmol^−1^Å^−2^ unit. RMSD(t) is the instantaneous RMSD with respect to the target structure, and RMSD_0_ (t) is a linearly decreasing targeted RMSD function where its slope is fixed by the total simulation time (10 ns) and, the initial and final RMSDs between starting and target structures. To check force constant and initial velocity dependencies, we varied force constant *k* as F0=200 and F1=300 and initial velocities as V0, V1 and V2, and found that force constant in that range does not show any significant change and also the simulation does not depend on the initial velocity (**Figure S3C**). For our purpose, we have set *k* = 200 kcalmol^−1^Å^−2^. All simulations (CMD and TMD) (**Table S7**) were carried out at constant pressure (1 bar) and temperature (NPT ensemble). The free energy, Δ*G*(t) = *E_MM_ +* Δ*G_solv_* (t) − *TS*(t), where *E_MM_* is the molecular mechanics energy from MD simulations, Δ*G_solv_* is the free energy of solvation obtained using implicit solvent model, Poisson-Boltzmann Surface Area (PBSA) method, *T* temperature in Kelvin, and entropy *S* calculated using Normal Mode Analysis (Nmode), was calculated for every snapshot using an AMBER (Ambertools version 14.0) module MM-PBSA-Nmode (Case *et al*. 2005). All the trajectory analyses were carried out using trajectory analysis software grcarma (Koukos & Glykos 2013), AMBER (Ambertools version 14.0) and VMD. For visualization purposes Pymol and VMD were used.

## Results

### A wide range of over-expression of Gal3p turns on the *GAL* switch in the absence of galactose

Before we set out to isolate mutations that stabilize Gal3p in inactive conformation, we employed two complementary plate assays (Kar *et al*. 2014) to demonstrate the range of over-expression of Gal3p that can cause constitutive expression of *GAL* genes. For this purpose, we used two plasmid constructs, wherein *GAL3* transcription is driven by *CYC1* promoter (**Figure S4A**) either from single-copy (*pCJM3*) or from multi-copy background (*pYJM3*). In one of the assays, we spotted tenfold serially diluted cell culture of transformants of *Sc385* (*gal3Δ* strain bearing *pCJM3* or *pYJM3* on a Ura d.o. medium (to maintain the plasmid) containing glycerol plus lactate (gly/lac) as the sole carbon source along with 2-deoxy-galatcose (2-DG) (See **Table S1** for genotype of strains and **Table S2** for features of plasmids). If the over-expression of Gal3p activates the *GAL* switch constitutively (i.e., in the absence of galactose), the transformants cannot grow in the above medium. This is because, the constitutively expressed galactokinase would convert 2-DG to 2-deoxy-galactose-1-phosphate (2-DG-1-P), which cannot be further catabolised. Accumulation of 2-DG-1-P causes toxicity thereby preventing the transformant from growing in gly/lac medium containing 2-DG. Transformants of *gal3□* bearing *pYJM3* do not grow on gly/lac plates containing 2-DG (**Figure 1A**) as demonstrated previously (Das Adhikari *et al*. 2014). In this study, we demonstrate that *gal3□* strain transformed with *pCJM3*, (single-copy Gal3p expression plasmid) is also unable to grow on gly/lac 2-DG plates (**Figure 1A**) as compared to the vector control (*pCJM*). The second assay is dependent upon the ability of the over-expressed Gal3p to activate the *HIS3* expression from a *P_GAL1_::HIS3* cassette integrated in a *ScRKK1* (*gal1*Δ*gal3*Δ*his3*Δ*P_GAL1_::HIS3*) strain. Unlike the *Sc385* (*gal3*Δ) strain used in the previous assay, *ScRKK1* strain lacks the alternative signal transducer *GAL1* and thus allows us to test the constitutive activation of *GAL* genes upon over-expression of Gal3p in a genetic context wherein only Gal3p over-expression activates the *GAL* switch. This strain transformed with *pCJM3 pYJM3* grow on gly/lac medium lacking histidine, while the strain bearing the control plasmids *pYJM* or *pCJM* do not (**Figure 1B**). In addition to reproducing our previous observation, we demonstrate that over-expression of Gal3p driven from *CYC1* promoter in a single-copy plasmid background also leads to the constitutive activation of *GAL* genes as determined by monitoring the activation of *GAL1* promoter (**Figure 1**).

**Figure 1.**
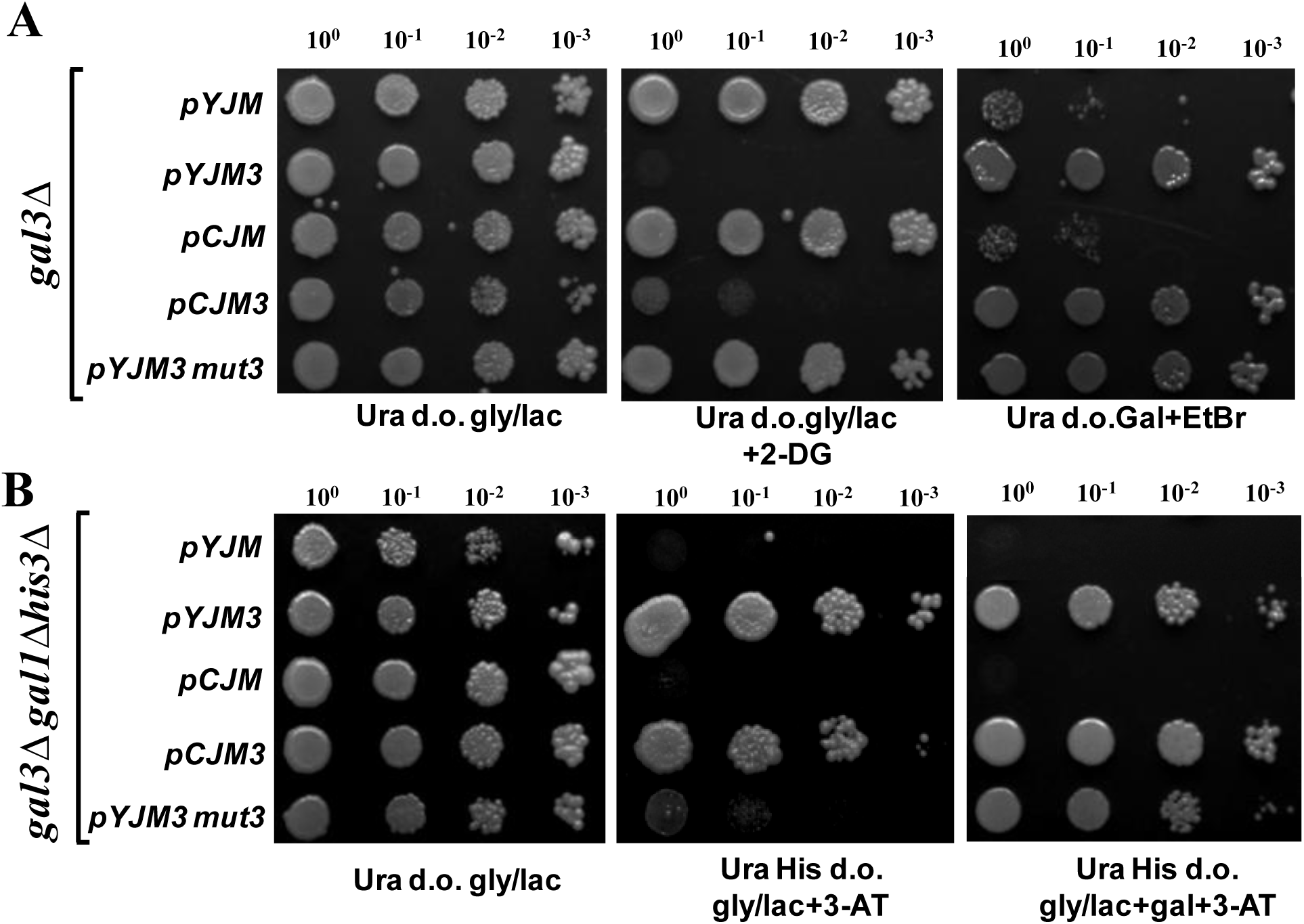
Over-expression of Gal3p driven by *CYC1* promoter from multi-copy (*pYJM3*) or single-copy plasmid (*pCJM3*) causes constitutive expression of *GAL* switch. **(A)** Overnight cultures of the transformants of *Sc385* (*gal3□* strain) bearing the indicated plasmids pre-grown on gly/lac medium were serially diluted and spotted onto URA drop out gly/lac plates containing 2-DG. EtBr was used to allow the growth of only those cells that utilize galactose as the sole carbon source and not the amino acids (Matsumoto *et al*. 1978). **(B)** Overnight cultures of the transformants of *ScRKK1* (*gal3*Δ*gal1* **Δ*his3*Δ*P_GAL1_****::His3*) bearing the indicated plasmids, pre grown on gly/lac medium were serially diluted and spotted onto the medium containing gly/lac as the sole carbon source but lacking histidine and uracil (His and Ura d.o.). *pYJM* and *pCJM* are the multi-copy and single-copy vector controls respectively. 3-amino-1,2,4-triazole (3-AT) reduces the background growth due to the leaky expression of *HIS3* (Blank *et al*. 1997).

### Isolation of inactive mutant of Gal3p, which is responsive to galactose

It was reported that the stability of mutant forms of Gal3p is less as compared to the wild-type (Platt *et al*. 2000). Based on this prior knowledge, we chose to introduce mutations in *GAL3* ORF in the multi-copy plasmid background, that is, in *pYJM3* (**Figure S4A**). The reason is that in case the mutations introduced causes reduced stability, then over-expression of mutant Gal3p’s from the multi-copy plasmid could compensate for the reduced stability and therefore be able to score the phenotype of the mutant Gal3p. We introduced random mutations using error prone PCR (EPPCR) in *GAL3* ORF in *pYJM3* (See **Table S3** and **S4** for EPPCR conditions) and recovered the mutant plasmid populations as URA^+^ transformants of *gal3*Δ strain (**Figure S4B**). These transformants were then replica plated on to Ura d.o. medium containing gly/lac plus 2-DG to score for 2-DG toxicity as well as galactose plates containing EtBr to score for growth on galactose. The transformants that grow on both the plates would contain the plasmids bearing the *GAL3* ORF with desired mutations.

The rational of the above protocol is as follows. Over-expression of those mutant Gal3p’s that remain in inactive conformation would not constitutively activate the expression of galactokinase and therefore would not cause toxicity. Such transformants would grow on gly/lac plate containing 2-DG. Any mutation that knocks off the function completely would also be picked up as a false positive in this screen. However, such mutants would not grow on the plates containing galactose plus EtBr and hence can be easily weeded out (**Figure S4B**). Using this dual screening strategy, we initially isolated ~ 50 transformants (Out of total 50,000 transformants) that showed the expected phenotype under both the assay conditions. Plasmids were isolated from these yeast transformants, amplified in *E.coli* and retransformed into the parent strain to confirm the phenotype. After going through this exercise several times to weed out the false positive plasmids, we finally obtained three putative Gal3p mutant plasmid clones, *pYJM3mut1, pYJM3mut2* and *pYJM3mut3*. By employing a plasmid loss experiment strategy, we confirmed that the phenotype is a plasmid borne trait and not due to genomic mutations (data not shown). We also determined the identity of the plasmids bearing the Gal3p mutant ORF, using restriction digestion analysis.

The above three putative plasmids were then subjected to 2-DG toxicity assay and His growth assay. As expected, independent transformants of *gal3*Δ strain bearing *pYJM3mut1, pYJM3mut2* and *pYJM3mut3* grew on gly/lac 2-DG plates, indicating that the above three plasmids were unable to confer constitutive expression of *GAL1* (**Figure S5A**). These transformants grew on EtBr plates suggesting that they responded to galactose activation. However, when these three plasmids were tested in His growth assay by transforming *ScRKK1* (*gal1* Δ*gal3* Δ*his3* Δ*P_GAL1_::HIS3*) strain, *pYJM3mut1* and *pYJM3mut2* were able to cause constitutive expression of *P_GAL1_::HIS3*, which is not expected, while *pYJM3mut3* failed to confer constitutive expression as expected (**Figure S5B**). Thus, *pYJM3mut3* does not confer constitutive expression of *GAL* genes when tested under two different experimental conditions, while *pYJM3mutl* and*pYJM3mut2* did not yield the expected phenotype. Since *pYJMmut1* and *2* failed to give the consistent phenotype in both the assays, we did not analyze these plasmids further. The inability of *pYJMmut3* plasmid to confer constitutive expression of *GAL* genes with appropriate controls is shown (**Figure 1**).

### Over-expression of mutant Gal3p does not result in the constitutive activation of *GAL* genes

Since constitutive activation of *GAL* genes (i.e.in the absence of galatcose) is dependent on the Gal3p concentration, it was important to establish the relative levels of wild-type Gal3p expression driven by *CYC1* promoter from a multi-copy (*pYJM3*) and single-copy plasmids (*pCJM3*) as well as the mutant Gal3p expressed from multi-copy plasmids (*pYJM3Mut3*). To determine this, western blot analysis was performed and the bands were empirically quantitated (see **Materials and Methods** and **Figure S6**), based on the dilution of the protein samples and using the loading control as the reference. Gal3p was found to be approximately 90 times more in cell free extracts obtained from the transformant bearing multi-copy plasmid (*pYJM3*) as compared to the single-copy plasmid (*pCJM3*) (**Figure S6**). Although the amount of Gal3p expressed from single-copy plasmid (*pCJM3*) was less by approximately 90 times as compared to the protein expressed from multi-copy (*pYJM3*), it was sufficient to give a clear cut read out of constitutive expression using two independent plate assays (**Figure 1**). Western blot analysis indicates that the steady state level of mutant Gal3p (*pYJM3Mut3*) was at least 10 times more than what is produced from the single-copy background (*pCJM3*) (**Figure S6**). That is, wild-type Gal3p expressed from single-copy (*pCJM3*), which is 10 times less than the mutant Gal3p (*pYJM3Mut3*) activates *GAL1* constitutively, as demonstrated by the plate assay (**Figure 1**). Based on these results, we conclude that the mutant Gal3p is far less efficient than the wild-type protein in constitutively activating the switch.

### Three spatially well-separated substitutions are necessary to maintain Gal3p in inactive and galactose responsive conformation

The *GAL3* ORF present in *pYJM3mut3* plasmid has V273I, T404A and N450D substitutions (**Figure S7**, see **Materials and Methods**). To determine whether any of the single or double mutations in any of the three combinations is sufficient to confer the unique property described above, we generated the following plasmids with single (*pYJM3*^V273I^*, pYJM3*^T404A^ and *pYJM3*^N450D^), or double mutations in all the three combinations (*pYJM3*^IA^, *pYJM3*^AD^ and *pYJM3*^DI^) and a triple mutation (*pYJM3*^IAD^) using SDM, for further analysis (See **Table S5** for primers for SDM in *GAL3* ORF). The transformants bearing these mutant plasmids were subjected to the plate assay as described previously. First, the transformant bearing *pYJM3mut3* (original isolate, see **Figure 1**) and *pYJM3*^IAD^ (generated using SDM) exhibited the same phenotype indicating that the phenotype is due to the mutations in *GAL3* ORF and not due to mutations elsewhere in the plasmids (**Figure 1 and 2**). Second, transformants bearing any of the single mutations, *pYJM3*^I^*, pYJM3*^A^ and *pYJM3*^D^ as well as double mutations, *pYJM3*^IA^, and *pYJM3*^DI^ express *GAL1* constitutively (**Figure 2**). This indicates that these mutants Gal3p’s behave more like the wild-type Gal3p. However, transformant bearing *pYJM3*^AD^ was unable to constitutively activate the switch but was not responsive to galactose unlike the *pYJM3*^IAD^.

**Figure 2.**
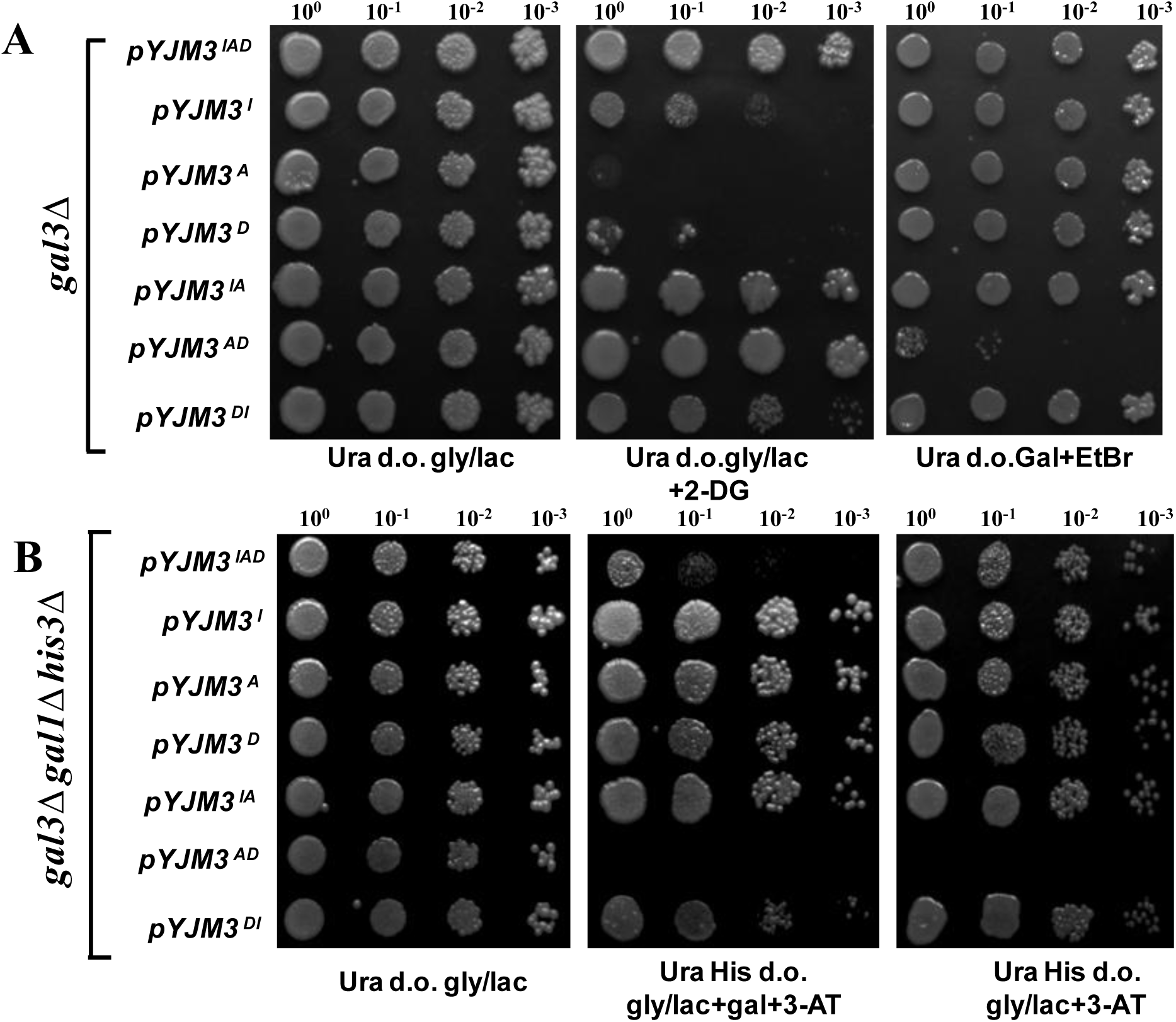
Three mutations are required to confer the inactive conformation of Gal3p. Transformants of **(A)** *Sc385* (*gal3*Δ) or **(B)** *ScRKK1* (*gal3*Δ*gal1*Δ*his3*Δ*P_GAL1_::His3*) strain bearing *pYJM3^lAD^* (triple mutant generated using site directed mutagenesis approach), *pYJM3*^I^*, pYJM3*^A^, pYJM3^D^(single mutants), *pYJM3*^IA^*, pYJM3*^AD^*, pYJM3*^DI^ (double mutants) pre-grown in gly/lac were serially diluted and subjected to **(A)** 2-DG toxicity assay and **(B)**. His growth assay.

Quantification of the western blot analysis using densitometric scanning revealed that the expression of Gal3p^IAD^ and Gal3p^AD^ were comparable (**Figure 3A**) while the Gal3p^IAD^ was at least 5 times more than the WT Gal3p expressed from single-copy (*pCJM3*). The above data indicate that Gal3p^AD^ is inactive and not responsive to galactose, while Gal3p^IAD^ is inactive but responsive to galactose (**Figure 2**). Thus, Gal3p^IAD^ has a higher propensity than the wild-type Gal3p to exist in inactive conformation. In addition to the phenotypic analysis (**Figure 2**), the *P_GAL1_::GFP* expression was monitored in *ScRKK39* (*gal3*Δ*gal1*Δ*P_GAL1_::GFP*) strain transformed with the plasmids containing WT Gal3p from single-copy (*pCJM3*), Gal3p^IAD^ and Gal3p^AD^ expressed from multi-copy plasmid and the corresponding vector (*pYJM*) (**Figure 3B**). The *P_GAL1_::GFP* expression was monitored in the transformants grown in gly/lac, in the absence of galactose (reflects constitutive activation of the *GAL* switch) and in the presence of galactose (reflects the responsiveness of Gal3p towards galactose). The mean fluorescence intensity of gly/lac grown *pYJM* (vector control), *pCJM3, pYJM3*^IAD^ and *pYJM3*^AD^ transformants are 54.4, 76.96, 48.41 and 68.41, respectively. The mean fluorescence intensity of gly/lac + galactose grown transformants are 56.84, 422.41, 376.04, and 63.55 (**Table S6**). As a control, we also monitored the constitutive (gly/lac grown) *P_GaL1_::GFP* expression in the transformants bearing *pYJM3* (multi-copy plasmid carrying wild-type *GAL3*) and *pYJM* (multi-copy vector control). As expected, the transformants carrying *pYJM3* growing in gly/lac had higher GFP expression than its corresponding vector control (**Figure S8** and **Table S6**).

**Figure 3.**
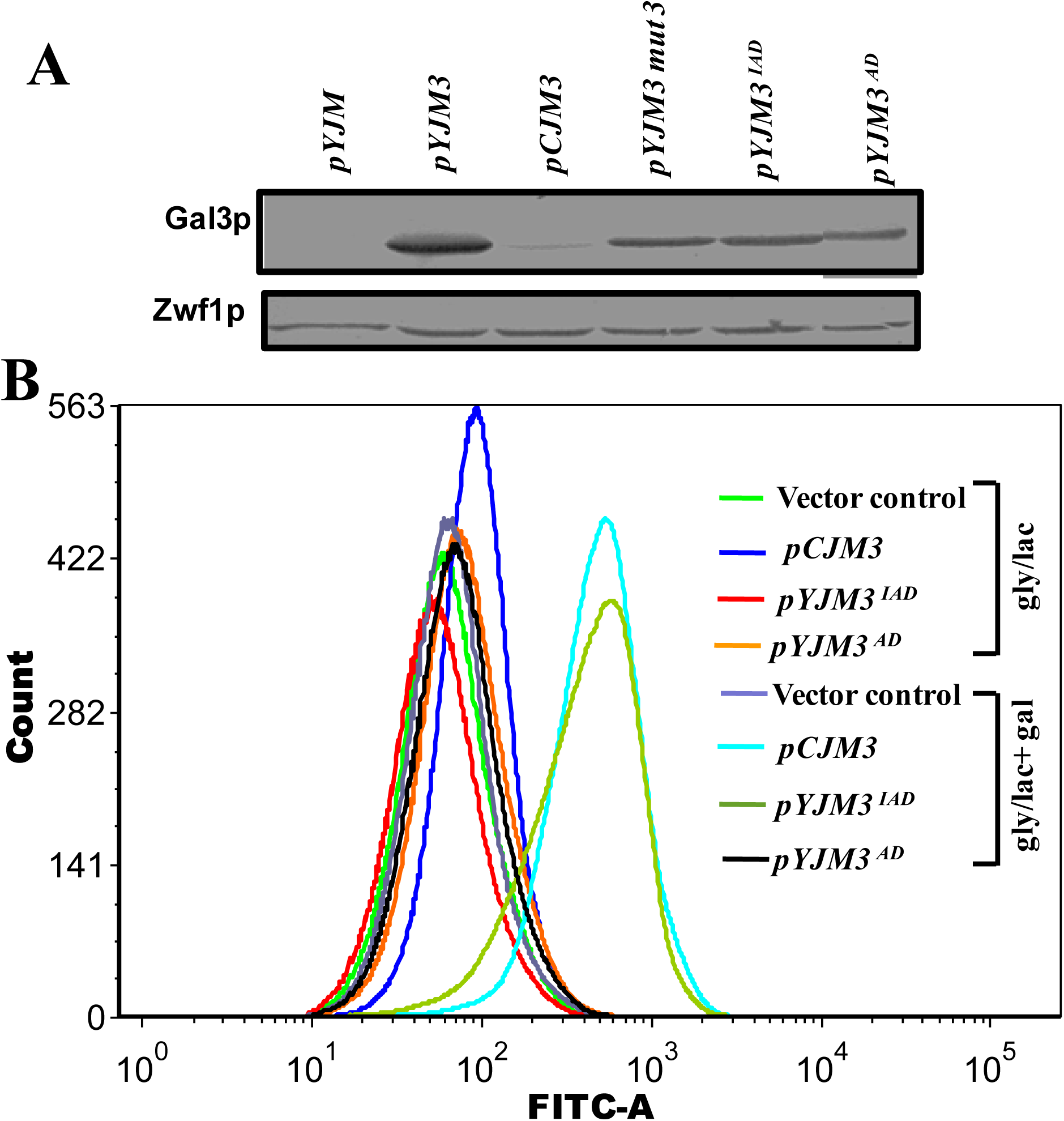
Western blot analysis of Gal3ps and constitutive expression of the *GAL* switch. **(A)** Cell free extracts obtained from gly/lac grown *ScRKK5* (*gal1*Δ*gal3*Δ) transformant bearing *pYJM* (vector control) *pYJM3, pCJM3, pYJM3^1AD^* (triple mutant generated using site directed mutagenesis approach), *pYJM^AD^* (double mutants) were subjected to western blot analysis. Zwf1p is used as the loading control. **(B)** Transformants of *ScRKK39* (*gal1*Δ*gal3*Δ*P_GAL1_::GFP*) bearing *pYJM* (vector control), *pCJM3* (Gal3p over expressed from single-copy plasmid), *pYJM3^1AD^* (Gal3p triple mutant generated using site directed mutagenesis approach over expressed from multi-copy plasmid) and *pYJM3^AD^* (Gal3p double mutants over expressed from multi-copy plasmid) were grown in gly/lac (shown in bright green, blue, red and orange color respectively) and gly/lac+galatcose (shown in blue gray, turquoise, lime and black color respectively) till mid-log. 50,000 cells were subjected to FACS analysis as described in material and methods. The mean fluorescence intensity of GFP was measured in FITC channel of flow cytometer. For the detail results of the FACS analysis refer **Table S6**. The experiment is repeated at least thrice with different set voltage and all the three patterns were same, therefore one set of data collected at a particular voltage is overlaid in this figure for comparison. (**Figure 3B** is a part of **figure S8** and is shown for clarity in comparing the overlaid mean GFP intensity of indicated plasmid transformant).

Thus, above results indicate that the constitutive *PGAL1*::*GFP* expression is lower in *pYJM3*^IAD^ (Gal3p^IAD^) as compared to *pCJM3* (**Figure 3B**) even though the protein expression is 10 times more than the wild type Gal3p (**Figure 3A**) as confirmed from the western blot analysis. Like wild type Gal3p, in response to galactose, Gal3p^IAD^ activates the switch, thus suggesting that it predominantly exists in inactive conformation. Similar to Gal3p^IAD^, Gal3p^AD^ is also unable to activate the switch constitutively. However, unlike Gal3p^IAD^, it is not responsive to galactose (See **Table S6** for statistical parameters of FACS analysis). Overall, these results suggest that all the three substitutions (V273I, T404A and N450D) are essential to stabilize the inactive conformation without sacrificing the ability of Gal3p^IAD^ to respond to galactose. We carried out molecular dynamics analysis of wild type and mutant forms of Gal3p to further elucidate the possible mechanism of the conformational transition.

### Closed states of Gal3p show conformational bimodality

The crystal structure of Gal3p in closed conformation shows a lip-distance of ≈ 30 Å between N-domain lip (D100) and C-domain lip (V375) (**Figure S9**). In the open conformation, these lips open up to ≈ 40 Å. It has been suggested that the opening and closing movement between these lips is related to the protein function (Menezes *et al*. 2003; Lavy *et al*. 2012). To understand the role of amino acid substitutions in the conformational dynamics of the protein, we have performed canonical molecular dynamics (CMD) simulations on both closed and open conformations of WT Gal3p and mutants of Gal3p. In **Figure 4**, the distance between residues D100 and V375 as a function of time (**Figure 4A and C**) and the corresponding conformational distributions (**Figure 4B and D**) are shown.

**Figure 4:**
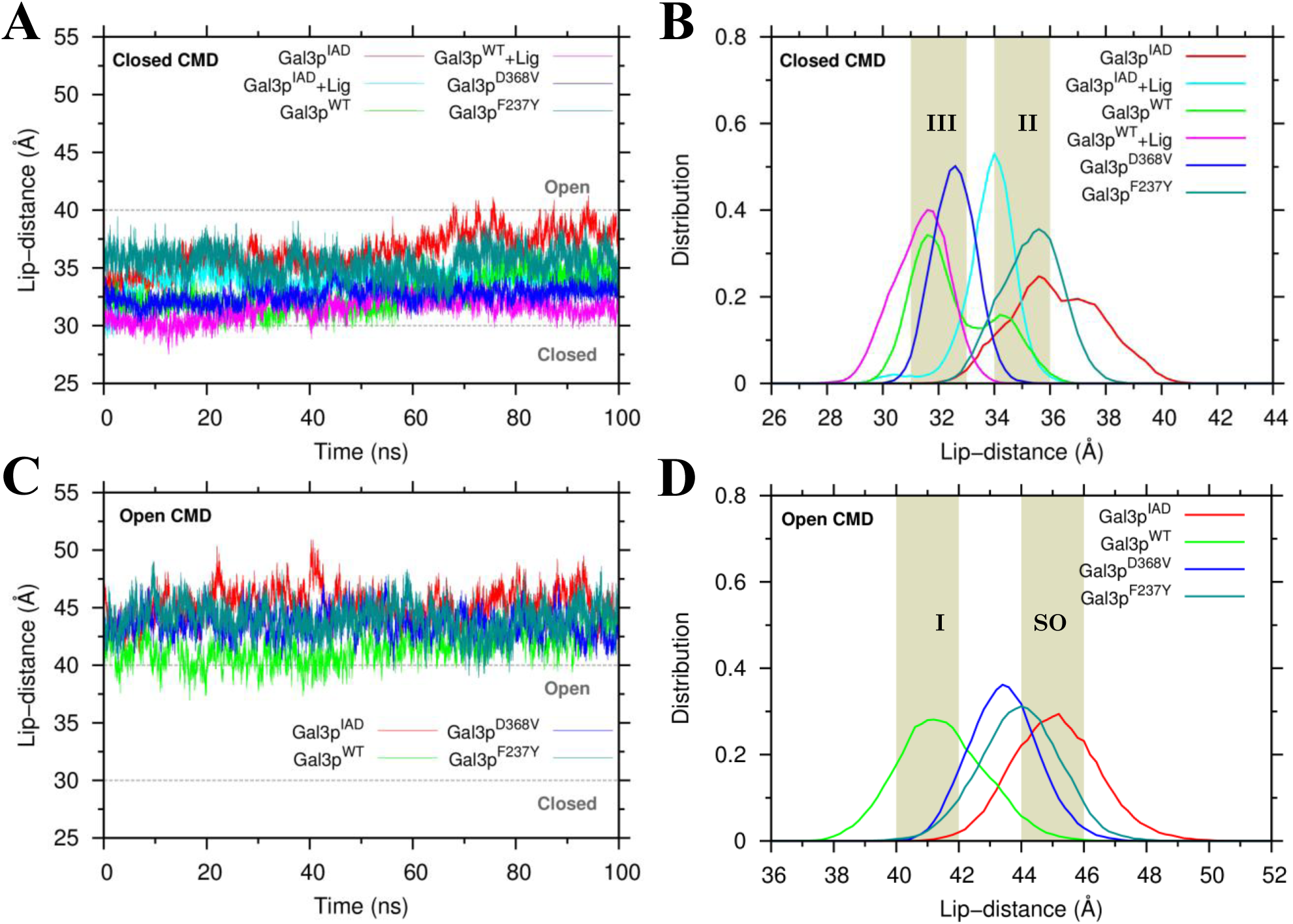
Conformational equilibrium dynamics. **(A, C)** show time progression of distance between N-domain lip (residue 100) and C-domain lip (residue 375) and **(B, D)** show their corresponding distributions, respectively. **(A, B)** were simulated starting from closed conformation while **(C, D)** were simulated using open conformation as the starting structure. The grey dash lines in **A** and **C** indicate the corresponding distances found in X-ray crystal structures.

In CMD starting with the closed structure (**Figure 4A and B**), the apo-forms of Gal3p^WT^ (WT Gal3p) and Gal3p^IAD^ (mutant of Gal3p with V273I, T404A, N450D substitutions) display contrasting dynamics with respect to the inter-domain motion. The lip-distance for Gal3p^WT^ shows a bimodal distribution with one peak lying in the region ≈ 31-33 Å and the other peak in the region ≈ 34-36 Å. In contrast, Gal3p^IAD^ is characterized by a broad distribution (spanning a region of ≈ 35-41 Å) suggesting that the apo-form of Gal3p^IAD^ has the propensity to adopt open conformations. However, in the presence of ligands the dynamics shift towards closed conformation. The liganded form of WT (Gal3p^WT^+Lig) shows a single peak. Interestingly, Gal3p^IAD^ with ligand (Gal3p^IAD^ +Lig), on the other hand, behaves like the apo-form of Gal3p^WT^, showing bimodal distribution with similar nature of peak locations.

We also simulated the behaviour of the previously isolated constitutive mutants (Gal3p^D368V^ and Gal3p^F237Y^) which do not require any ligand for activation of *GAL* genes (Blank *et al*. 1997). Both Gal3p^WT^+Lig and Gal3p^D368V^ have the same range of lip-distance ≈ 31-33 Å. In the case of Gal3p^F237Y^, its highest peak falls in the region ≈ 34-36 Å (**Figure 4A and B**). Therefore, we surmise that conformations with lip-distance ≈ 31-33 Å and 34-36 Å are “Gal3p^D368V^-like” (III) and “Gal3p^F237Y^-like” (II) active conformations respectively, which bind to Gal80p in the absence of ligands.

In CMD starting with an open conformation (**Figure 4C and D**), all the variants show single peaks and interestingly all the mutants (Gal3p^IAD^, Gal3p^D368V^ and Gal3p^F237Y^) sample structures with higher lip-distance than Gal3p^WT^, especially Gal3p^IAD^ with lip-distance ≈ 44-46 Å as opposed to Gal3p^WT^ ≈ 40-42 Å. These structures are stable and well separated from the closed structures, an indication that CMD can hardly cross high energy barriers. However, by starting the simulation from different initial conformations, we can indeed obtain information about the local dynamics and important insights in understanding the experiments. We classify structures with lip-distance falling in the range ≈ 40-42 Å as “open” (I) and those falling in range ≈ 44-46 Å as “super-open” (SO). The justification for the conformational classification becomes apparent from the results discussed in the following section.

### Free energy of activation pathway reveals stable intermediate states

In order to access intermediate states and to complement our CMD results, we used targeted molecular dynamics (TMD) simulations and sampled all possible structures starting from open-state (lip-distance > 40 Å) to closed-state (lip-distance < 32 Å). To determine conformational stability, free energy (see computational details in **Materials and Methods**) was calculated at every time point. As the proteins undergo conformational transition along the activation pathway, the lip-distances continue to decrease until they reach ≈ 7 ns, the time point from where all the variants start to converge to ≈ 32-33 Å, except for Gal3p^F237Y^, which converges to ≈ 36 Å (**Figure 5A**). We call this region, which spans from 7 to 10 ns, the “region of stability” (shaded region, **Figure 5A**). Remarkably, in the region of stability, all the structures seem to assume single conformation with respect to lip orientation. We believe that using lip-distance as a large-scale conformational order-parameter, our TMD has been successful in scanning all the possible structures between open- and closed-states.

**Figure 5:**
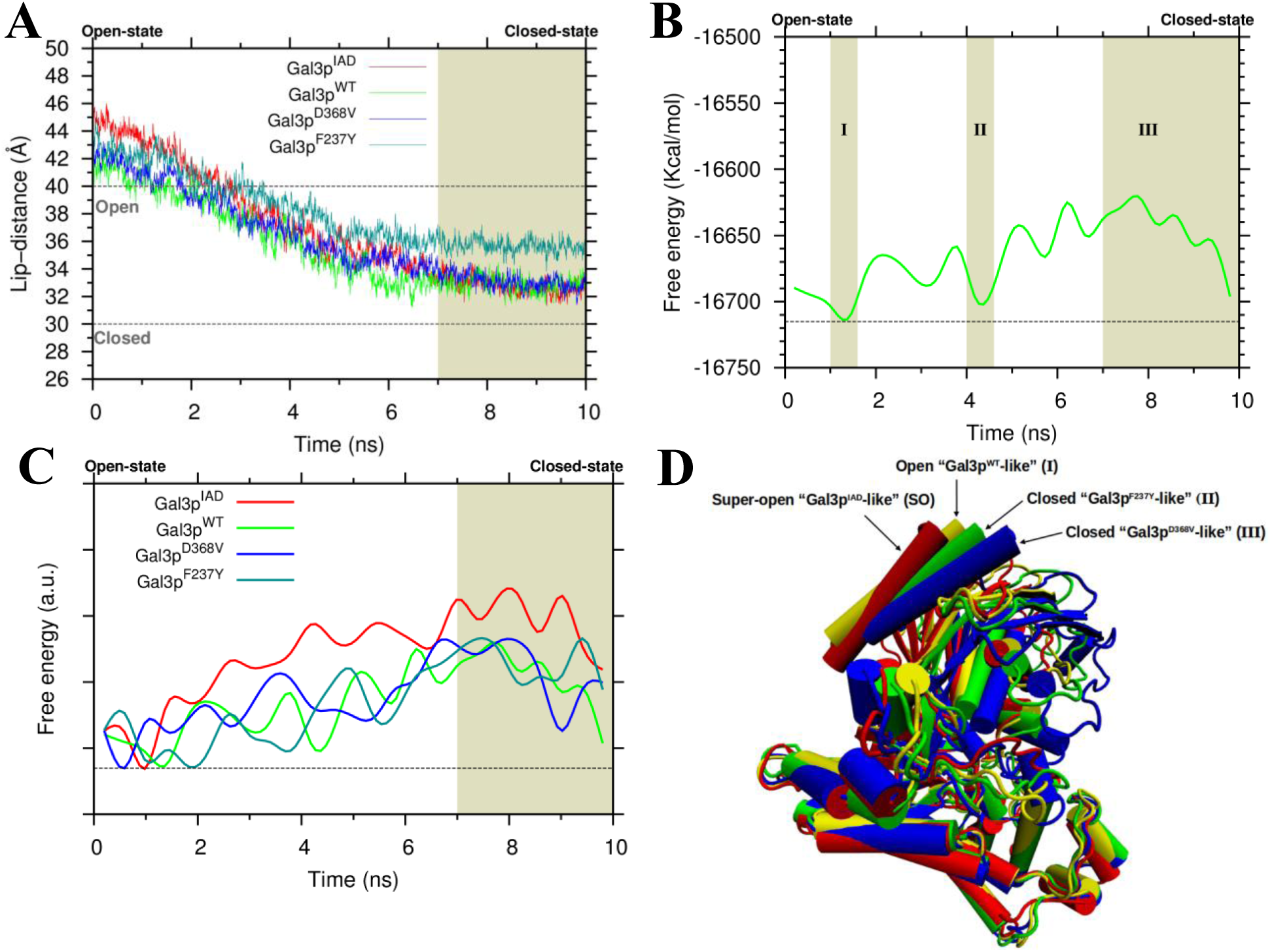
Allosteric activation pathway. **(A)** Time evolution of Lip-distance along activation pathway. The grey dash lines indicate the corresponding distances in closed and open crystal structures. (**B** and **C**) Conformational free energy along activation pathway. **(B)** The free energy landscape for Gal3p^WT^ and **(C)** shows plot for all the variants where the global minima are placed on the same energy level represented by black dash lines. The grey shaded regions indicate minima along the pathway and labeled I and II. **(A-C)** The region of stability is shown as shaded region in olive color and also labeled as III in **(B). (D)** Superposition of structural representatives from regions I (yellow), II (green), III (blue) in **(B)** and the global minimum of Gal3p^IAD^ (red) in **(C)** with respect to their C-domains: Conformation I, II and III are the open “Gal3p^WT^-like”, closed “Gal3p^F237Y^-like” and closed “Gal3p^D368V^-like” structures respectively, and the conformation shown in red is the super-open “Gal3p^IAD^-like” structure.

Free energy landscape along a potential transition pathway for the WT (Gal3p^WT^) is shown in **Figure 5B**. The result indicates that there are three important conformations: I (≈ 1 ns), II (≈ 4 ns) and III (≈ 7-10 ns). Conformation I corresponds to the global minimum with lip-distance ≈ 40-42 Å and it overlaps with the crystal structure of apo-form (open) of Gal3p^WT^. Conformation II represents a local minimum, which corresponds to a lip-distance of ≈ 34-36 Å and possibly a stable intermediate state. We predict that this intermediate state is “Gal3p^F237Y^ - like” active conformation. Structures lying in the olive shaded region (III) have lip-distance of ≈ 31-33 Å and correspond to higher energy states in the absence of ligands (compare conformations I, II and III in **Figure 5B**). We predict that this closed state is “Gal3p^D368V^-like” active conformation. Comparison of free energy landscapes for all the variants is shown in **Figure 5C**. It can be seen that the initial structures in the first nanosecond (TMD ≈ 1 ns) are their respective global minima (**Figure 5C**). As the TMD progresses along the activation pathway, the free energy increases for all the variants. Strikingly, all the variants except Gal3p^IAD^, show stable intermediate states (minima), whereas for Gal3p^IAD^, the intermediate states have higher energies. In the region of stability (shaded region, **Figure 5C**), Gal3p^IAD^ structures have higher energies compared to other variants, while Gal3p^D368V^ has a low stable minimum. Gal3p^WT^ and Gal3p^F237Y^, on the other hand, have intermediate energy levels. In **Figure 5D**, we present superposition of structures taken from regions I, II, III (**Figure 5B**), and the global minima of Gal3^IAD^ (**Figure 5C**), with respect to their C-domains. We found that between any consecutive structures there is a rotation of ≈ 15° in their N-domains. Thus, the lip opening in super-open state is 45° of rotation more than the closed state.

### Residues T404A and N450D stabilize inactive conformation while V273I stabilizes active conformation

It was previously demonstrated that F237, a hydrophobic hinge residue, plays a pivotal role in stabilising the closed conformation of Gal3p^F237Y^ (Lavy *et al*. 2012). In the WT protein, F237 is seated in the “hydrophobic pocket” formed by residues F247, M403, L65, F414, F418, and I245 (**Figure 6D**; the hydrophobic pocket is indicated in grey color). It was proposed that the spatial orientation of this residue directly governs the structural change of the protein, with a solvent-exposed side-chain resulting in a closed conformation and its burial within the hydrophobic pocket resulting in an open conformation (Lavy *et al*. 2012). In this study, solvent accessible surface area (SASA) of residue 237 with respect to time was determined to study the effects of mutations on its orientation (**Figure 6A**-**C**). It is clearly seen from the results that, in closed conformer dynamics, residue F237 in Gal3^IAD^ tends to remain buried, whereas in the Gal3p^WT^, Gal3p^D368V^ and Gal3p^F237Y^, it prefers to pop out into the solvent (**Figure 6A**). However, in the open conformer dynamics, all the variants have residue 237 buried almost all the time except for Gal3p^F237Y^, which tends to remain exposed to solvent (**Figure 6B**). The histogram of SASA of this residue also demonstrates that open (and super-open), intermediate and closed conformations have SASA ≈ 15 Å^2^, 30 Å^2^ and 50-60 Å^2^ respectively (**Figure S10**). SASA from TMD convincingly tells us that F237 in Gal3p^IAD^ prefers to remain buried (**Figure 6C**). We suggest that the T404A substitution, which lies close to the hydrophobic pocket, extends the hydrophobic pocket and thus allows the F237 to get stably buried in Gal3p^IAD^ (**Figure 6D**). Additionally, the salt-bridge formation between residue Asp at position 450 (N450D) and residue Lys at position 488, constrains the relative motion between helices a15 and a16 (**Figure 6D and E**). Thus, mutant of Gal3p with both the substitutions is unable to activate the switch even in the presence of galactose (**Figure 2A and B**).

**Figure 6:**
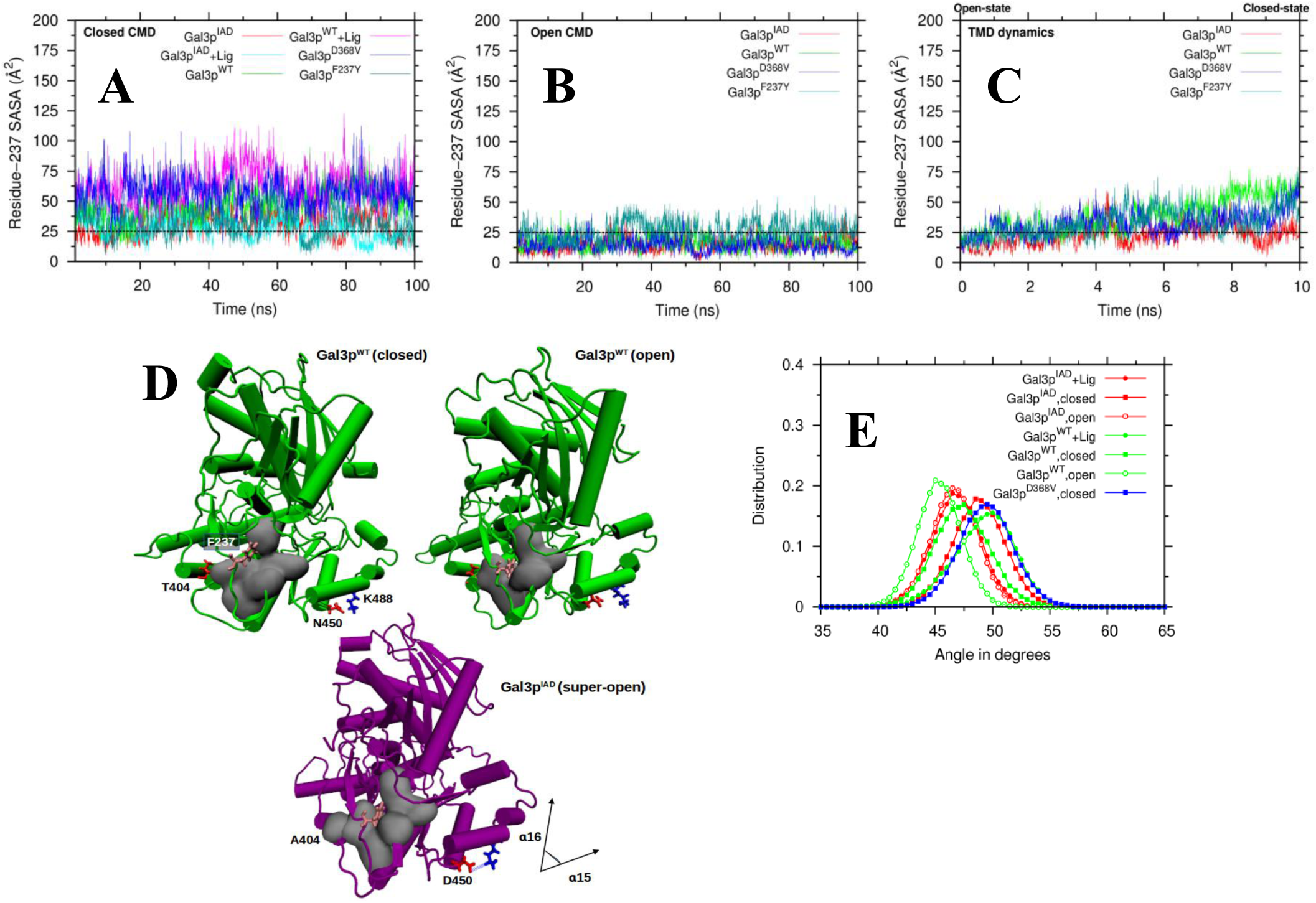
Role of T404A and N450D in stabilization of inactive conformations. SASA of residue 237 with respect to time is presented in (**A-C**). (**A**) and (**B**) represent equilibrium dynamics (CMD) and (**C**) represents biased dynamics (TMD). Since calculation of SASA of any residue from interior protein is < 25 Å^2^, SASA ≤ 25 Å^2^ is considered buried, whereas SASA > 25 Å^2^ is considered exposed to the solvent (black dash line). **(D)** Snapshots of CMD structures of Gal3p^WT^ (green), in apo-closed (average SASA ≈ 50 Å^2^) and open (16 Å^2^) states, and Gal3p^IAD^ (purple) in open (15 Å^2^) state, showing different orientations of residue F237 (pink) seated in the hydrophobic pocket formed by residues F247, M403, L65, F414, F418, and I245 (silver) in the hinge region. In Gal3p^IAD^, mutations T404A (red) forms extended hydrophobic pocket and mutations N450D (red) forms salt-bridge with residue K488 (blue). **(E)** Distributions of angle between helices α15 and α16.

Root mean square deviations (RMSD) from TMD clearly show that both N- and C-domains undergo significant local rearrangement with C-domain having higher degree of rearrangement as compared to the N-domain along the activation pathway (**Figure 7A and B**). In all the protein variants, the nature of RMSD evolution in N-domain is more or less the same. Strikingly, the RMSD of C-domain for Gal3p^IAD^ is the highest and remain constant from ≈ 7 ns onwards (**Figure 7A** shaded portion), suggesting the stabilizing role of V273I substitution when the protein adopts active (closed) conformation. One of the major contributions to structural rearrangement in C-domain may have come from the “polar-loop” fluctuation (**Figure 7C** and **D**). This is evident from the RMSF calculation (**Figure S11A** and **S12**) showing that there is higher fluctuation in open state than in closed state. In open state, this loop becomes more solvent accessible and its conformational entropy increases. This conformational entropy is minimized in closed state when the two lips come closer. This is important because residues 298 and 299 in the polar-loop are involved in interaction with Gal80p. Furthermore, the residue 299 forms a salt-bridge with residue 368 in all the protein variants, except for Gal3p^D368V^, irrespective of whether the global conformation is in closed or open state. Interestingly, in the closed state, the loop adopts certain orientation in order to accommodate special structure of the salt-bridge between 299 and 368 (**Figure 7E**; also see the snapshot at TMD = 8 ns). In Gal3p^D368V^, the residue 299 is perfectly trapped in the hydrophobic groove stabilized by D368V substitution in the same manner that reduces conformational entropy of the loop (**Figure 7F**).

**Figure 7:**
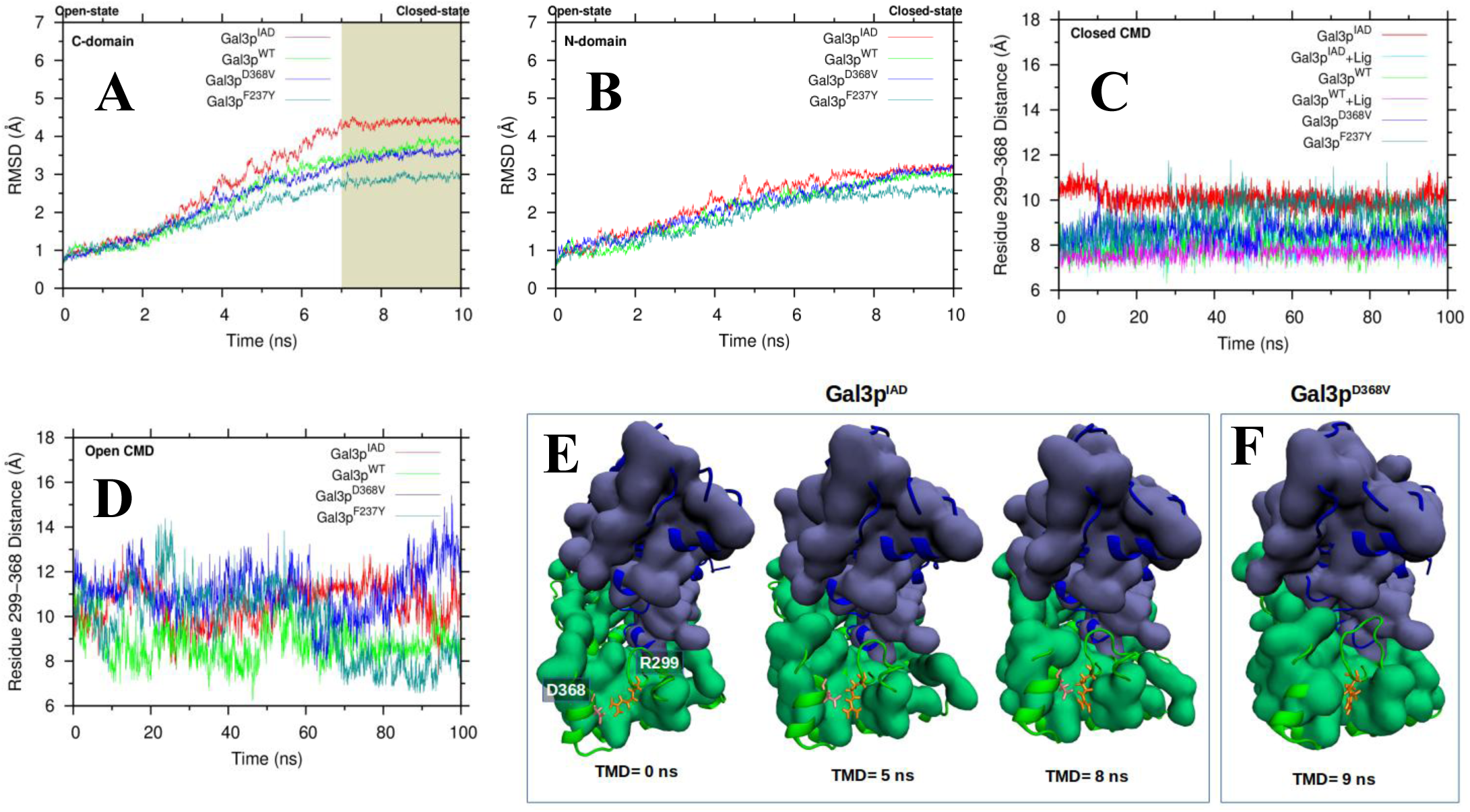
Role of V273I in stabilization of active conformations. (**A** and **B**) Domain-RMSD from TMD trajectory. (**C** and **D**) Polar-loop (LII) dynamics from CMD trajectory. Distance between residues 299 (from LII) and 368 are plotted as a function of time. (**E** and **F**) TMD snapshots at different time points. Hydrophobic residues are shown in surface rendition and the polar residues are shown in cartoon representation. N-domain is colored blue and C-domain in green. Residues 299 (orange) and 368 (pink) that forms salt-bridge in all the protein systems (closed dynamics and open dynamics) except in constitutive mutant, Gal3p^D368V^, are shown in licorice representation. (**E**) Various structural forms of salt-bridge interaction between D368 and R299. (**F**) Structural representative from a minimum lying in the olive-shaded region (**Figure 5C** blue curve). In this mutant, residue D368V forms a part of the stable hydrophobic C-domain surface.

## Discussion

While previous genetic and structural studies had suggested that wild-type Gal3p can exist in active and inactive conformations (Bhat & Hopper 1992; Lavy *et al*. 2012), the possible mechanism of allosteric transition was not deciphered. We decided to study the allosteric transition by using molecular dynamics simulation (MD), as Gal3p is too large to study using NMR. However, MD of only wild-type Gal3p and the constitutive mutants could be performed and thus, our access was limited to only a part of the spectrum of conformations that a wild-type Gal3p and its constitutive derivatives can potentially exist. Therefore, we were constrained by the non-availability of mutant derivatives of Gal3p that would stabilize the inactive conformation. To overcome this lacuna, we isolated Gal3p^IAD^ that appears to have high propensity to exist predominantly in inactive conformation, which is at the other end of the conformational spectrum. Using MD of mutant that we isolated (Gal3p^IAD^), we were able to capture a super-open (SO) conformation, in addition to I, II and III reported previously (Lavy *et al*. 2012). The molecular dynamic analysis of the Gal3p^IAD^ is in consonance with conclusions drawn based on the experimental results thus corroborating our molecular dynamic analysis. This work has led us to unravel many unknown facets of allosteric transitions in Gal3p.

Although the positions of the three mutations are located far apart from each other and away from the Gal80p interaction sites (**Figure S11B**), we can rationalize the importance of their strategic location in stabilizing the inactive conformation without losing its ability to interact with galactose. For example, the residue T404A lies adjacent to the hydrophobic pocket where it harbors the side chain of residue F237. Substitution of residue T by hydrophobic residue A at position 404 results in the formation of extended hydrophobic pocket, so that the side chain of F237 gets stably buried in the hydrophobic pocket of Gal3p^IAD^. Alteration of neutral charge residue N to negatively charge residue D at position 450 leads to salt-bridge formation between residue N450D and K488. An interesting consequence of such a perturbation in the local interaction environment is that, it affects the relative motion between helices a15 and a16 via conformational restriction (**Figure 6D and E**). Thus, it appears that the combined effect due to co-substitution of T404A and N450D in the background of V273I substitution provides structural stability to SO conformation in the absence of galactose. This could be attributed to the fact that, Gal3p^AD^ exists predominantly in region I and in the region between I and II (~36-38Å) (**Figure S13B, C** and **D**). Additionally, our MD and free energy calculations also reveal that Gal3p^AD^ also accesses region III (**Figure S13A** and **D**). Interestingly, Conformation III is immune to galactose since the closed state does not allow the ligand to bind. Conformations with lip-distance greater than 36Å may have low affinity for galactose. This may explain why Gal3p^AD^ does not response to galactose. Contrastingly, in Gal3p^ID^, the conformation with the minimum free energy corresponds to region II (**Figure S13A** and **D**). Moreover, in Gal3p^IA^ and Gal3p^ID^, V273I is a conservative substitution and thus may not contribute towards the stabilization of the SO state. These results demonstrate that only these constellations of substitutions, V273I, T404A and N450D, can give rise to the phenotype unique to Gal3p^IAD^.

We suggest that the reason for the loss of the ability of Gal3p^IAD^ to activate constitutively the *GAL* system is because, it exists mainly in SO conformation. Our analysis clearly indicates that Gal3p^WT^ visits the closed (III), intermediate (II), and open (I) states more often than the SO state. In other words, Gal3p^WT^ visits SO state very rarely. In contrast, in the Gal3p^IAD^ the SO structure is the predominant form. It is interesting to note that three independent mutations are essential to stabilize the SO structure that confers the unique property to the protein. This is in sharp contrast to the stabilization of the closed structure that can be brought about by introducing substitutions at single sites (Blank *et al*. 1997), which are spread across the primary sequence. It is possible that mutations that can give rise to constitutive phenotype could be obtained by destabilizing the inactive conformation as well. Unlike this, stabilizing the SO conformation need a specific constellation of mutations.

Our analysis can be extended to explain the previously isolated constitutive mutations. A mutation at the position 237 from F to Y (F237Y) confers the protein the ability to constitutively activate the *GAL* switch (Blank *et al*. 1997). The position of this residue is located at the hinge region and has been suggested that upon substitution of F by Y, the polar side chain of residue Y interacts with the hydrophilic solvent thereby facilitating the protein to adopt active state (Lavy *et al*. 2012). Consistent with this, we observe that the side chain of F237 remains buried in the open (I) and SO structures, while it remains exposed to the solvent in the closed structures. Thus the exposure of residue 237 to solvent is correlated with lip closure and vice versa. Our simulation results suggest that mutation F237Y stabilizes the active intermediate states (II) (**Figure 5B**).

Another constitutive mutation D368V located on the C-domain, which interacts with Gal80p (Blank *et al*. 1997), is an example of mutations lying far away from the hinge region (**Figure S11B**). Incidentally, the residue R299, which lies in the polar loop (LII) region (**Figure S9B**) forms a salt-bridge with D368 (**Figure 7E**). We have shown that lip open-closure movement and the structural local rearrangement due to salt-bridge interaction between residues D368 and R299 in the C-domain, are correlated. However, the salt-bridge that stabilizes the closed states (III) via reduction in fluctuations of polar loop (LII) region, is very different from the salt bridge found in open (I) and super-open (SO) structures. Therefore, we surmise that, the substitution of D by V, a hydrophobic residue, in D368V mutant, may stabilize the hydrophobic surface and allow residue 368 to be perfectly localized in the hydrophobic groove. We provide an explanation on the basis of structure and dynamics how mutation D368V stabilizes the closed state (III) (**Figure 5B**).

In the light of our results, we propose a dynamic conformational model for allosteric activation of Gal3p (**Figure 8**). The intrinsic fluctuation of F237 side chain has been shown to correlate with domain-domain (lip) motion. When analyzing tertiary H–bonds networks (excluding secondary structure H–bonds i.e. H–bonds involved in helices and sheets formation), we found that the inter-domain H–bonds are not as extensive as compared to the intra-domain H-bonds (**Figure S14**). This observation also supports the idea that the protein exhibits easy lip movement. Thus, the protein can be modeled as a “hinge-wedge-pacman model” where the residue F237 plays the role of a pivot-wedge at the hinge region. As shown in **Figure 8**, Gal3p may preexist in four states defined by the order-parameters: lip-distance between the domains and the SASA of F237. In the absence of galactose, wild-type Gal3p exists mostly in energetically favorable open state (I). However, when galactose is present, it promotes “conformational selection” on I, II and SO. We assume that galactose can bind to I, II and SO except state III, but with decreasing affinity as the lip-distance increases. We surmise that structure II has the highest affinity for galactose and once bound, this conformation may undergo “induced-fit” to adopt more thermodynamically stable conformation III (closed state). The closed structures, on the other hand, may be stabilized by ligand-mediated H-bonds and/or hydrophobic interactions (residues: P112, V114, V201, V203, F259, A262, P263 and F373; (See **movie S1**)). Therefore, we predict that the probable activation pathway may consist of I, II, III and SO traced out on the parameter map defined by lip-distance and SASA of F237.

**Figure 8:**
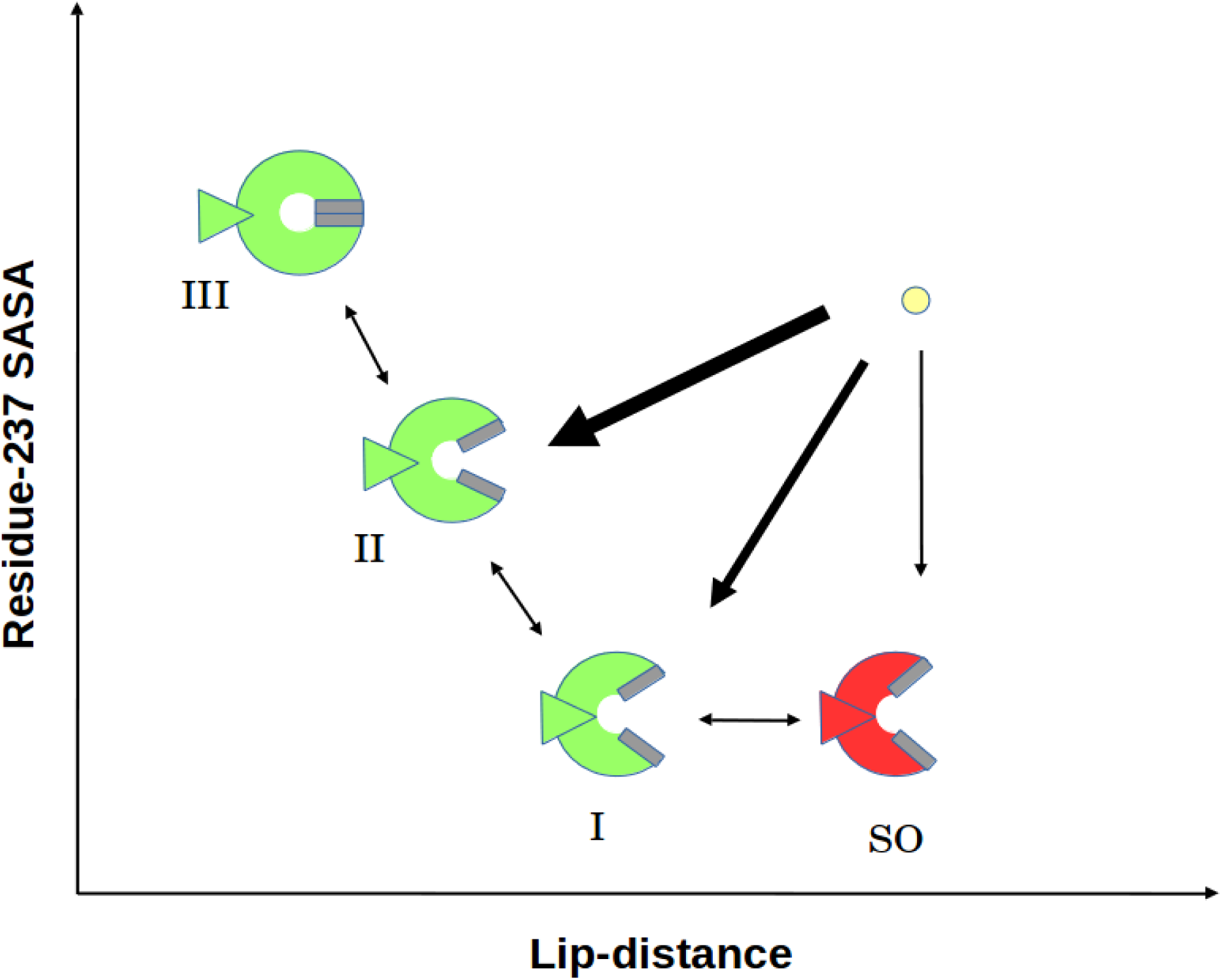
Allosteric conformational model for Gal3p. Gal3p is represented as “hinge-wedge-pacman” model where the hinge-wedge (triangle) depicts the residue F237 and the hydrophobic lips (grey boxes) represent the residues: 112, 114, 201, 203, 259, 262, 263 and 373. Gal3p pre-exists in four conformations: I, II and III (green) are energetically favourable structures and SO (red) is energetically less favourable structure. The presence of galactose (yellow circle) induces “conformational selection” on I, II and SO structures and encourages “induced-fit” between II and III structures. We propose that galactose binds to II with high affinity and binds to SO with low affinity (indicated by thickness of the arrows).

Human glucokinase, a monomeric bifunctional protein, is involved in the catalysis of glucose phosphorylation as well as signal transduction. Similar to Gal3p, its interaction with a downstream target protein is an integral part of signal transduction process (de, I *et al*. 2000). Structural study has demonstrated that it exists predominantly in super-open (SO) state in the absence of glucose (Kamata *et al*. 2004). Conformational restriction to open-closed states by a small allosteric compound alters substrate saturation curve from an inherent “sigmoidal” to acquired “hyperbolic” pattern (Kamata *et al*. 2004). Such a large-scale conformational transition between SO and closed states with stable intermediate states has been shown to be responsible to sigmoidal pattern of substrate saturation curve. Similar to what we observe in Gal3p, the allosertic site in glucokinase also consists of a hydrophobic pocket (HP) (Huang *et al*. 2013) made up of Tyr162, Val200, Ala201, Met202, Val203, Leu5451, Val42, Cal455 and Ala 456 with Ile 159, which is analogous to F237 of Gal3p. As discussed before, a similar HP exists in Gal3p, where the side chain of residue F237 is docked. Our results have demonstrated that solvent accessibility of this side chain is correlated with lip motion. In closed state, HP is found to be “inside-out” making it unfavorable for docking which causes residue 237 side chain to remain exposed to the solvent. Whereas, in the open (super-open) state, the HP forms a deep pocket favorable for docking.

Galactokinase of *S. cerevisiae* is a paraloge of Gal3p and can also transduce the galactose induced signal of the *GAL* genetic switch. Galactokinase activity shows a hyperbolic saturation pattern with respect to galactose (Timson & Reece 2002). These proteins share a high sequence similarity, so much so, that corresponding residues V273, T404 and N450 of Gal3p are conserved in Gal1p (**Figure S7**). If SO structure is responsible for sigmoidicity in glucokinase, we predict that substitutions at equivalent positions by I, A and D should convert Gal1p from a hyperbolic to a sigmoidal protein. It is intriguing to note that although glucokinase (hexokinase family) and Gal3p (GHMP family) belong to different families, they seem to follow similar mechanism of conformational transitions suggesting that such mechanisms are conserved and could be found in other proteins as well.

To the best of our knowledge, there are two reports of the isolation of triple mutant that confer a unique phenotype (Iiri *et al*. 1999; Alper *et al*. 2006). Of the two, the more relevant to our study is the triple mutant of G□ mutant A366S, G226A and E268A(Iiri *et al*. 1999), which dominantly inhibits the receptor activation. None of the single or double mutation in any of the combinations exhibits the negative dominant property. Interestingly, these mutations are highly conserved and one of them is involved in the elimination of salt bridge. Salt bridges have been shown to be responsible for conformational stability regardless of where they exist in the overall protein structure (Kumar & Nussinov 1999; Bosshard *et al*. 2004). It appears that salt bridges in general seem to have a vital role in allosteric transition. Thus, obtaining a unique conformation with which a specific property is associated (in this case, Gal3p^IAD^ is inactive but responsive to galactose) provides us a handle to study allostery resulting from protein dynamics. This attempt highlights the importance of isolating mutations with unique properties and subjecting them to detailed molecular dynamics simulation for elucidating the structural basis of allosteric mechanisms.

## Acknowledgements

We thank CRNTS of IITB for flow cytometry facility. We acknowledge the financial support from IITB, CSIR, DST and 08BRNS004. We thank Dr. R. Komel for gifting *YCpGAL1GFP* plasmid. We also thank YUVA PARAM supercomputing facilities, Pune, for MD simulations purposes.

## Author Contributions

Wet lab experiments were conceived and designed by PJB and executed by RKK. MD simulations were mainly conceived and performed by HK with input from RP. The paper was mainly written by PJB and HK wrote all of the MD part. RP contributed in refining the discussion. Performed all the experiments: RKK. Performed all the molecular dynamics simulation: HK. Analyzed the data: RKK HK RP PJB. Contributed reagents/materials/analysis tools: RP PJB.

## Literature Cited

Alper, H., J. Moxley, E. Nevoigt, G. R. Fink, and G. Stephanopoulos, 2006 Engineering yeast transcription machinery for improved ethanol tolerance and production. Science 314: 1565–1568.

Bhat, P. J., and J. E. Hopper, 1992 Overproduction of the GAL1 or GAL3 protein causes galactose-independent activation of the GAL4 protein: evidence for a new model of induction for the yeast GAL/MEL regulon. Mol. Cell Biol. 12: 2701–2707.

Blacklock, K., and G. M. Verkhivker, 2014 Computational modeling of allosteric regulation in the hsp90 chaperones: a statistical ensemble analysis of protein structure networks and allosteric communications. PLoS. Comput. Biol. 10: e1003679.

Blank, T. E., M. P. Woods, C. M. Lebo, P. Xin, and J. E. Hopper, 1997 Novel Gal3 proteins showing altered Gal80p binding cause constitutive transcription of Gal4p-activated genes in Saccharomyces cerevisiae. Mol. Cell Biol. 17: 2566–2575.

Boehr, D. D., R. Nussinov, and P. E. Wright, 2009 The role of dynamic conformational ensembles in biomolecular recognition. Nat. Chem. Biol. 5: 789–796.

Bosshard, H. R., D. N. Marti, and I. Jelesarov, 2004 Protein stabilization by salt bridges: concepts, experimental approaches and clarification of some misunderstandings. J. Mol. Recognit. 17: 1–16.

Brooks, B. R., C. L. Brooks, III, A. D. Mackerell, Jr., L. Nilsson, R. J. Petrella et al. 2009 CHARMM: the biomolecular simulation program. J. Comput. Chem. 30: 1545–1614.

Case, D. A., T. E. Cheatham, III, T. Darden, H. Gohlke, R. Luo et al. 2005 The Amber biomolecular simulation programs. J. Comput. Chem. 26: 1668–1688.

Cooper, A., and D. T. Dryden, 1984 Allostery without conformational change. A plausible model. Eur. Biophys. J. 11: 103–109.

Das Adhikari, A. K., M. T. Qureshi, R. K. Kar, and P. J. Bhat, 2014 Perturbation of the interaction between Gal4p and Gal80p of the Saccharomyces cerevisiae GAL switch results in altered responses to galactose and glucose. Mol. Microbiol. 94: 202–217.

de, l., I, M. Mukhtar, J. Seoane, J. J. Guinovart, and L. Agius, 2000 The role of the regulatory protein of glucokinase in the glucose sensory mechanism of the hepatocyte. J. Biol. Chem. 275: 10597–10603.

Gietz, R. D., and R. A. Woods, 2002 Transformation of yeast by lithium acetate/single-stranded carrier DNA/polyethylene glycol method. Methods Enzymol. 350: 87–96.

Gunasekaran, K., B. Ma, and R. Nussinov, 2004 Is allostery an intrinsic property of all dynamic proteins? Proteins 57: 433–443.

Hammes, G. G., Y. C. Chang, and T. G. Oas, 2009 Conformational selection or induced fit: a flux description of reaction mechanism. Proc. Natl. Acad. Sci. U. S. A 106: 13737–13741.

Huang, M., S. Lu, T. Shi, Y. Zhao, Y. Chen et al. 2013 Conformational transition pathway in the activation process of allosteric glucokinase. PLoS. One. 8: e55857.

Humphrey, W., A. Dalke, and K. Schulten, 1996 VMD: visual molecular dynamics. J. Mol. Graph. 14: 33–38.

Iiri, T., S. M. Bell, T. J. Baranski, T. Fujita, and H. R. Bourne, 1999 A Gsalpha mutant designed to inhibit receptor signaling through Gs. Proc. Natl. Acad. Sci. U. S. A 96: 499–504.

James, L. C., P. Roversi, and D. S. Tawfik, 2003 Antibody multispecificity mediated by conformational diversity. Science 299: 1362–1367.

James, L. C., and D. S. Tawfik, 2003 Conformational diversity and protein evolution-a 60-year-old hypothesis revisited. Trends Biochem. Sci. 28: 361–368.

Kamata, K., M. Mitsuya, T. Nishimura, J. Eiki, and Y. Nagata, 2004 Structural basis for allosteric regulation of the monomeric allosteric enzyme human glucokinase. Structure. 12: 429–438.

Kar, R. K., M. T. Qureshi, A. K. Dasadhikari, T. Zahir, K. V. Venkatesh et al. 2014 Stochastic galactokinase expression underlies GAL gene induction in a GAL3 mutant of Saccharomyces cerevisiae. FEBS J. 281: 1798–1817.

Kern, D., B. F. Volkman, P. Luginbuhl, M. J. Nohaile, S. Kustu et al. 1999 Structure of a transiently phosphorylated switch in bacterial signal transduction. Nature 402: 894–898.

Kjelsberg, M. A., S. Cotecchia, J. Ostrowski, M. G. Caron, and R. J. Lefkowitz, 1992 Constitutive activation of the alpha 1B-adrenergic receptor by all amino acid substitutions at a single site. Evidence for a region which constrains receptor activation. J. Biol. Chem. 267: 1430–1433.

Koshland, D. E., 1958 Application of a Theory of Enzyme Specificity to Protein Synthesis. Proc. Natl. Acad. Sci. U. S. A 44: 98–104.

Koukos, P. I., and N. M. Glykos, 2013 Grcarma: A fully automated task-oriented interface for the analysis of molecular dynamics trajectories. J. Comput. Chem. 34: 2310–2312.

Kumar, S., and R. Nussinov, 1999 Salt bridge stability in monomeric proteins. J. Mol. Biol. 293: 1241–1255.

Laemmli, U. K., 1970 Cleavage of structural proteins during the assembly of the head of bacteriophage T4. Nature 227: 680–685.

Lavy, T., P. R. Kumar, H. He, and L. Joshua-Tor, 2012 The Gal3p transducer of the GAL regulon interacts with the Gal80p repressor in its ligand-induced closed conformation. Genes Dev. 26: 294–303.

Lin-Goerke, J. L., D. J. Robbins, and J. D. Burczak, 1997 PCR-based random mutagenesis using manganese and reduced dNTP concentration. Biotechniques 23: 409–412.

Lockless, S. W., and R. Ranganathan, 1999 Evolutionarily conserved pathways of energetic connectivity in protein families. Science 286: 295–299.

Marsh, J. A., and S. A. Teichmann, 2015 Structure, dynamics, assembly, and evolution of protein complexes. Annu. Rev. Biochem. 84: 551–575.

Matsumoto, K., Toh-e A, and Y. Oshima, 1978 Genetic control of galactokinase synthesis in Saccharomyces cerevisiae: evidence for constitutive expression of the positive regulatory gene gal4. J. Bacteriol. 134: 446–457.

McDonald, L. R., J. A. Boyer, and A. L. Lee, 2012 Segmental motions, not a two-state concerted switch, underlie allostery in CheY. Structure. 20: 1363–1373.

Menezes, R. A., C. Amuel, R. Engels, U. Gengenbacher, J. Labahn et al. 2003 Sites for interaction between Gal80p and Gal1p in Kluyveromyces lactis: structural model of galactokinase based on homology to the GHMP protein family. J. Mol. Biol. 333: 479–492.

Monod, J., J. Wyman, and J. P. Changeux, 1965 ON THE NATURE OF ALLOSTERIC TRANSITIONS: A PLAUSIBLE MODEL. J. Mol. Biol. 12: 88–118.

Motlagh, H. N., and V. J. Hilser, 2012 Agonism/antagonism switching in allosteric ensembles. Proc. Natl. Acad. Sci. U. S. A 109: 4134–4139.

Murthy, T. V., and B. P. Jayadeva, 2000 Disruption of galactokinase signature sequence in gal3p of Saccharomyces cerevisiae does not lead to loss of signal transduction function. Biochem. Biophys. Res. Commun. 273: 824–828.

Okazaki, K., and S. Takada, 2008 Dynamic energy landscape view of coupled binding and protein conformational change: induced-fit versus population-shift mechanisms. Proc. Natl. Acad. Sci. U. S. A 105: 11182–11187.

Phillips, J. C., R. Braun, W. Wang, J. Gumbart, E. Tajkhorshid et al. 2005 Scalable molecular dynamics with NAMD. J. Comput. Chem. 26: 1781–1802.

Platt, A., H. C. Ross, S. Hankin, and R. J. Reece, 2000 The insertion of two amino acids into a transcriptional inducer converts it into a galactokinase. Proc. Natl. Acad. Sci. U. S. A 97: 3154–3159.

Platt, T., 1984 Toxicity of 2-deoxygalactose to Saccharomyces cerevisiae cells constitutively synthesizing galactose-metabolizing enzymes. Mol. Cell Biol. 4: 994–996.

Sali, A., L. Potterton, F. Yuan, V. H. van, and M. Karplus, 1995 Evaluation of comparative protein modeling by MODELLER. Proteins 23: 318–326.

Sambrook J.F.E.F.a.M.T. Molecular cloning: a laboratory manual. (2^nd^ edition). 1989. Newyork, Cold Spring Harbor laboratory press, Cold Spring Harbor. Ref Type: Serial (Book, Monograph)

Schneider, C. A., W. S. Rasband, and K. W. Eliceiri, 2012 NIH Image to ImageJ: 25 years of image analysis. Nat. Methods 9: 671–675.

Schuster, M., R. E. Silversmith, and R. B. Bourret, 2001 Conformational coupling in the chemotaxis response regulator CheY. Proc. Natl. Acad. Sci. U. S. A 98: 6003–6008.

Silva, D. A., G. R. Bowman, A. Sosa-Peinado, and X. Huang, 2011 A role for both conformational selection and induced fit in ligand binding by the LAO protein. PLoS. Comput. Biol. 7: e1002054.

Sinha, N., and R. Nussinov, 2001 Point mutations and sequence variability in proteins: redistributions of preexisting populations. Proc. Natl. Acad. Sci. U. S. A 98: 3139–3144.

Stagoj, M. N., A. Comino, and R. Komel, 2005 Fluorescence based assay of GAL system in yeast Saccharomyces cerevisiae. FEMS Microbiol. Lett. 244: 105–110.

Timson, D. J., and R. J. Reece, 2002 Kinetic analysis of yeast galactokinase: implications for transcriptional activation of the GAL genes. Biochimie 84: 265–272.

Tsai, C. J., and R. Nussinov, 2014 A unified view of “how allostery works”. PLoS. Comput. Biol. 10:e1003394.

Vogt, A. D., and C. E. Di, 2012 Conformational selection or induced fit? A critical appraisal of the kinetic mechanism. Biochemistry 51: 5894–5902.

Volkman, B. F., D. Lipson, D. E. Wemmer, and D. Kern, 2001 Two-state allosteric behavior in a single-domain signaling protein. Science 291: 2429–2433.

Weber, G., 1972 Ligand binding and internal equilibria in proteins. Biochemistry 11: 864–878.

William L. Jorgensen J.C.J.D.M.R.W.I.a.M.L.K. Comparison of simple potential functions for simulating liquid water. J. Chem. Phys. 79[3], 926–935. 1983. Ref Type: Journal (Full)

Zhang, J., C. Li, K. Chen, W. Zhu, X. Shen et al. 2006 Conformational transition pathway in the allosteric process of human glucokinase. Proc. Natl. Acad. Sci. U. S. A 103: 13368–13373.

